# Modelling tick bite risk by combining random forests and count data regression models

**DOI:** 10.1101/642728

**Authors:** Irene Garcia-Marti, Raul Zurita-Milla, Arno Swart

## Abstract

The socio-economic and demographic changes occurred over the past 50 years have dramatically expanded urban areas around the globe, thus bringing urban settlers in closer contact with nature. Ticks have trespassed the limits of forests and grasslands to start inhabiting green spaces within metropolitan areas. Hence, the transmission of pathogens causing tick-borne diseases is an important threat to public health. Using volunteered tick bite reports collected by two Dutch initiatives, here we present a method to model tick bite risk using human exposure and tick hazard predictors. Our method represents a step forward in risk modelling, since we combine a well-known ensemble learning method, Random Forest, with four count data models of the (zero-inflated) Poisson family. This combination allows us to better model the disproportions inherent in the volunteered tick bite reports.

Unlike canonical machine learning models, our method can capture the overdispersion or zero-inflation inherent in data, thus yielding tick bite risk predictions that resemble the original signal captured by volunteers. Mapping model predictions enables a visual inspection of the spatial patterns of tick bite risk in the Netherlands. The Veluwe national park and the Utrechtse Heuvelrug forest, which are large forest-urban interfaces with several cities, are areas with high tick bite risk. This is expected, since these are popular places for recreation and tick activity is high in forests. However, our model can also predict high risk in less-intensively visited recreational areas, such as the patchy forests in the northeast of the country, the natural areas along the coastline, or some of the Frisian Islands. Our model could help public health specialists to design mitigation strategies for tick-borne diseases, and to target risky areas with awareness and prevention campaigns.

## 1 Background

Over the last couple of decades, urban areas have dramatically expanded (EEA, 2006). In Europe, the development of low density residential areas in the periphery of cities has become the norm for urban growth. (EEA, 2006). This phenomenon, known as urban sprawl, has a plethora of negative effects over the local climate (e.g. urban heat islands), the modification of the landscape (e.g. fragmentation), and the alteration of ecosystems (e.g. biodiversity loss) (EEA, 2011). In addition, urban sprawl also brings urban settlers in closer contact with nature and the countryside (Tack, 2013). As a response, several bird (e.g. thrushes) and mammal species (e.g. rodents, foxes, raccoons) have adapted their ethology to be able to live at the interface between forests and urban regions (e.g. more food, less predators) (Uspensky, 2017). This also means that the parasites and pathogens that several wildlife species carry get closer to residential areas and that, for instance, the hazard for tick-borne diseases increases (Allan, Keesing, & Ostfeld, 2003). In parallel to this, the progressive adoption of healthier lifestyles encourages citizens to spend more time outdoors carrying out leisure (Mulder, van Vliet, Bron, Gassner, & Takken, 2013) or sportive activities (Hall, Alpers, Bown, Martin, & Birtles, 2017). This behavior leads to a higher exposure to tick-borne diseases (Sandifer, Sutton-grier, & Ward, 2015).

Socio-economic changes and the subsequent response of nature means that citizens are more vulnerable to tick-borne diseases today than in the past. Hence, the transmission of pathogens causing tick-borne diseases is an important public health threat (Ehrmann et al., 2017). In fact, recent research demonstrates that ticks have trespassed the limits of forests and natural grasslands to start inhabiting green spaces within metropolitan areas. Urban parks in Zurich (Oechslin et al., 2017), Milan (Olivieri, Gazzonis, Zanzani, Veronesi, & Manfredi, 2017), Kiev (Didyk et al., 2017), Warsaw (Kowalec et al., 2017) or Lisbon (Santos et al., 2018), and suburban forests in Paris (Paul, Cote, Le Naour, & Bonnet, 2016), Budapest (Szekeres et al., 2016, 2018) or Wroclaw (Kiewra, Stefańska-Krzaczek, Szymanowski, & Szczepańska, 2017), present tick populations, and researchers were able to identify pathogens capable of causing Lyme borreliosis (LB) or tick-borne encephalitis (TBE) in humans. Since parks and suburban forests are potentially visited by thousands to millions of citizens every year, it is necessary to fully comprehend and model the risk of getting a tick bite to prevent this major public health threat.

The risk of getting a tick bite is the result of the interaction between its exposure and hazard components. Traditionally, researchers have tried to represent the risk of LB by quantifying the hazard component alone (Eisen et al., 2010; Gassner, Hansford, & Medlock, 2016; LoGiudice, Ostfeld, Schmidt, & Keesing, 2003), but in the last years researchers have worked on integrating hazard and exposure metrics to model tick bite risk (De Keukeleire et al., 2015; Zeimes, Olsson, Hjertqvist, & Vanwambeke, 2014) because hazard maps alone are insufficient to identify locations with a high risk for LB (Garcia-Marti, Zurita-Milla, Harms, & Swart, 2018).

The location of citizens is key to model the level of risk they are exposed to, but acquiring this type of information requires a partnership of researchers and public-health specialists to create (inter-)national networks of surveillance and citizen observatories. In the Netherlands, Wageningen University & Research (WUR) and the Dutch Institute for Public Health and the Environment (RIVM) started two citizen science projects to collect data on ticks and tick bites. These projects, which started in 2006 and 2012, have attracted enough media attention over the years to engage citizens at contributing tick bite reports. This engagement has resulted in over 50,000 volunteered tick bite reports in the Netherlands. This unique dataset enables new approaches to monitor and model elusive public health threats, such as tick bites. However, volunteered data is often unstructured, contains positional inaccuracies and reporting bias, and observations have a variable quality (Mehdipoor, Zurita-Milla, Augustijn, & van Vliet, 2018; Senaratne, Mobasheri, Ali, Capineri, & Haklay, 2017), conditions that might cause difficulties when including volunteered data in a scientific workflow.

A major challenge in our work was dealing with the highly skewed and zero-inflated distribution of the tick bite reports. These two types of disproportions are inherent traits of our data collection. Skewness refers to the asymmetry of the distribution, whereas zero-inflation refers to distributions in which zero-valued observations are frequent. In this work, our goal is creating a spatial tick bite risk model with national coverage. However, adding the spatial dimension implies the simultaneous modelling of a few locations reporting a high number of tick bites, and a substantial amount of locations in which zero tick bites are recorded.

Although these characteristics make it hard to use machine learning methods (Krawczyk, 2016), we pursue a solution based on machine learning because of its proven ability to handle non-linear and complex relationships. Classical count data statistical models are better equipped to handle skewed and zero-inflated datasets but they are unable to capture the inherent non-linearity in data. Thus, here we propose a solution integrating machine learning and classical statistic models. We combine the “segmentation capabilities” of the well-known Random Forest regressor (Breiman, 2001), which can split the data into homogenous groups using decision tree rules, with count data models of the Poisson family, which are better suited to model count data. Thus, our scientific contribution is two-fold: we present a tick bite risk model based on a wide array of hazard and exposure metrics, and we propose a methodological step forward by combining Random Forest and count data models to better model skewed and zero-inflated distributions.

## 2 Risk, exposure, and hazard

In the field of risk assessment, risk (R) is often modelled as a function of hazard (H) and exposure (E). The relationship between these three variables can be conceptualized as R = H × E (Braks, van Wieren, Takken, & Sprong, 2016). Thus, the calculation of tick bite risk should include variables representing both the H and E components, which likely have different underlying factors.

In the case of ticks and LB, the first spirochetes were identified in the early 1980s (Burgdorfer et al., 1982), and it took several years for the first studies to point out at human exposure to ticks as the source of the disease. Back then, various studies (e.g. Falco & Fish, 1989; Magnarelli, Denicola, Stafford, & Anderson, 1995) had already identified urban recreational parks as risky locations for LB, thus recommending prevention campaigns at parks and to inform citizens living nearby a green space. LB emerges from a complex ecological system driven by a wide array of factors (e.g. wildlife, climate, vegetation) (Ostfeld, 2012). For over 20 years scientists have studied the interactions between these factors to devise robust models of tick hazard. Multiple efforts can be found in literature since the late 1990s to quantify and map this component of tick bite risk. However, in our recent research (Garcia-Marti et al., 2018) we found out that the E component may be more relevant to determine tick bite risk. The quantification of the E component is a challenging task, due to the unavailability of human exposure metrics at the national scale. Thus, in this work we devoted special effort and creative thinking at developing novel human exposure indicators, which are combined with our tick hazard model (Garcia-Martí, Zurita-Milla, van Vliet, & Takken, 2017) to predict tick bite risk.

### 2.1 Tick hazard

The H component of tick bite risk has been widely studied since the late 1990s. Scientists have thoroughly worked to understand the impact that wildlife (Ostfeld, Canham, Oggenfuss, Winchcombe, & Keesing, 2006; Randolph & Storey, 1999), mast years (Buonaccorsi et al., 2003; Kelly, Koenig, & Liebhold, 2008), vegetation type (Tack, Madder, Baeten, De Frenne, & Verheyen, 2012), and weather variables (Berger, Ginsberg, Gonzalez, & Mather, 2014) have on tick populations. The pursuit of reliable models on tick hazard has prompted researchers to model this component of risk from multiple perspectives. Thus, we can find studies dedicated to tick habitat suitability (Estrada-Peña, de la Fuente, Latapia, & Ortega, 2015), tick presence (Swart et al., 2014), tick activity (Bennet, Halling, & Berglund, 2006), or tick dynamics (Garcia-Martí et al., 2017), with a varying number of biotic and abiotic parameters, and applied from local to continental spatial scales. In this work we use tick activity as a proxy for tick hazard. Tick activity represents the number of ticks that are questing for blood meals, which are the ones biting humans. Tick activity is extracted from a data-driven model that predicts daily tick activity in forests and natural grasslands (Garcia-Martí et al., 2017). The map in Figure 3 shows the predicted tick activity of this model, which is the average number of questing ticks per grid cell for the entire study period (2006-2014).

### 2.2 Human exposure

Human exposure to ticks is intrinsically linked to human behavior outdoors and to diverse socio-economic factors. For instance, (Zeman & Benes, 2014) discuss the peri-urbanization process in the Czech Republic, which prompted wealthy settlers to move away from large metropolitan areas into peri-urban areas to be in closer contact with nature and open spaces. This, in turn, triggered an increase in the number of tick-borne infections that was not directly related with any identifiable expansion on the tick habitat range.

Similarly, the societal adoption of healthier lifestyles encourages citizens to spend more time outdoors carrying out physical or sportive activities. For instance, in (Hall et al., 2017) the authors used a mass-participation cross-country marathon competition in Ireland to survey a large number of citizens and assess their exposure to ticks. Also, (Padgett & Bonilla, 2011) identify common picnic spots in a national park in the USA, as locations posing a risk of human exposure to ticks. Children participating in scouting or summer camp activities are found to be vulnerable to tick bites in a study in Belgium (De Keukeleire et al., 2015). All these examples are associated to the so-called “recreational exposure”, however, there are also studies that pinpoint activities in the peri domestic environment as risky for tick bites. Previous works considering the “residential exposure” include a study in the Netherlands finding a high risk of tick bites in gardens (Mulder et al., 2013) and two studies in the USA (Hahn et al., 2017) and Czech Republic (Zeman, Benes, & Markvart, 2015) indicating that properties in the peri-urban environment with a large interface between a forest and the garden or backyard pose a high risk for inhabitants of acquiring pathogens.

A thorough study for TBE in Stockholm demonstrates a use case in which exposure and hazard variables are combined to obtain tick bite risk indicators (Zeimes et al., 2014). The authors create metrics for human exposure based in accessibility and scenic beauty, whereas for the hazard they include variables of wildlife, forest, and land cover. Indeed, accessibility measured as the distance to an access road or trail is an important variable to account for when modelling tick bite risk. In (Li, Colson, Lejeune, & Vanwambeke, 2016) the authors assess the willingness of residents to travel for woodland leisure, because it varies as a function of whether citizens have to walk, cycle, or drive to the leisure place. However, accessibility is not the only factor to account for human exposure. There are locations in nature that are attractive for citizens due to presence of recreational areas or amenities (Lambin, Tran, Vanwambeke, Linard, & Soti, 2010), or because they have intrinsic value such as high scenic beauty or a good preservation (Nielsen, Heyman, & Richnau, 2012).

## 3 Methods

### 3.1 Tick bite risk monitored by volunteers

This work is based on the collection of volunteered tick bite reports from the Natuurkalender (NK; “nature’s calendar”; http://www.natuurkalender.nl) and the Tekenradar (TR; “tick radar”; https://www.tekenradar.nl) citizen science initiatives promoted by WUR and the RIVM. During the six years (2006-2012) that NK registered tick bite reports, this platform gathered 9,256 user contributions, whereas the TR initiative collected 46,655 reports for the period 2012-2016. This means that in total there are 55,911 reports available. However, some of these reports lack geographic coordinates or are placed outside the boundaries of the Netherlands. Hence, a total of 46,838 valid reports were found. Here we approximate the risk of getting a tick bite in a given area by the cumulative sum of tick bites reports in that area for the whole study period (i.e. 2006-2016).

Prior to the modelling phase a spatial aggregation operation was used to transform the individual tick bite reports into a tick bite risk proxy. We choose a regular grid with cells of 1km^2^ for the aggregation because that is the resolution of the existing hazard model described in Section 2.1. This aggregation groups together observations that are close in the geographic space (Figure 1a). However, it also creates a grid with right-skewed and zero-inflated (Figure 2) grid cell values. More precisely, the grid has a total of 36,866 cells. 9,985 cells have at least one tick bite report and the remaining 26,881 cells have zero tick bite reports. This means that for each grid cell with at least 1 tick bite, we have 3 in which no tick bites are recorded. Skewedness and zero-inflation are common real-world problems, especially when modelling count data, (Hadiji, Molina, Natarajan, & Kersting, 2015). Thus the analysis of the tick bite reports requires a modelling approach capable of handling these characteristics (Krawczyk, 2016).

**Figure 1.**
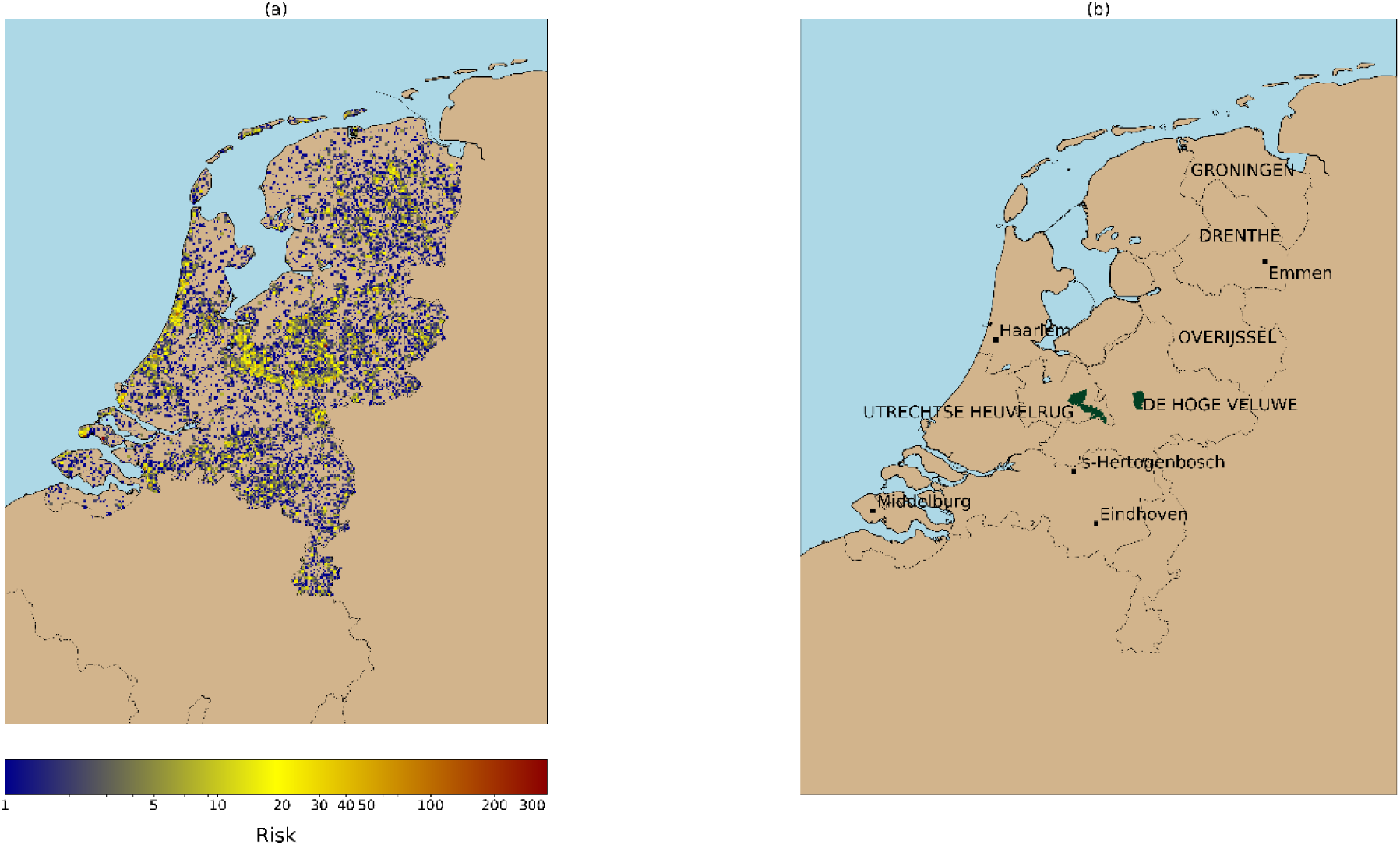
(a) Tick bite risk (2006-2016) as monitored by the NK and TR initiatives. We use as a proxy of tick bite risk the cumulative sum of tick bite reports per 1km grid cell. This image shows that tick bites are produced throughout the country, but the reports tend to be clustered around forests (e.g. Veluwe national park), or recreational areas (e.g. coastal areas). (b) Geographic locations. Provinces and national parks are labeled with capital letters, cities are labeled in lowercase.

**Figure 2.**
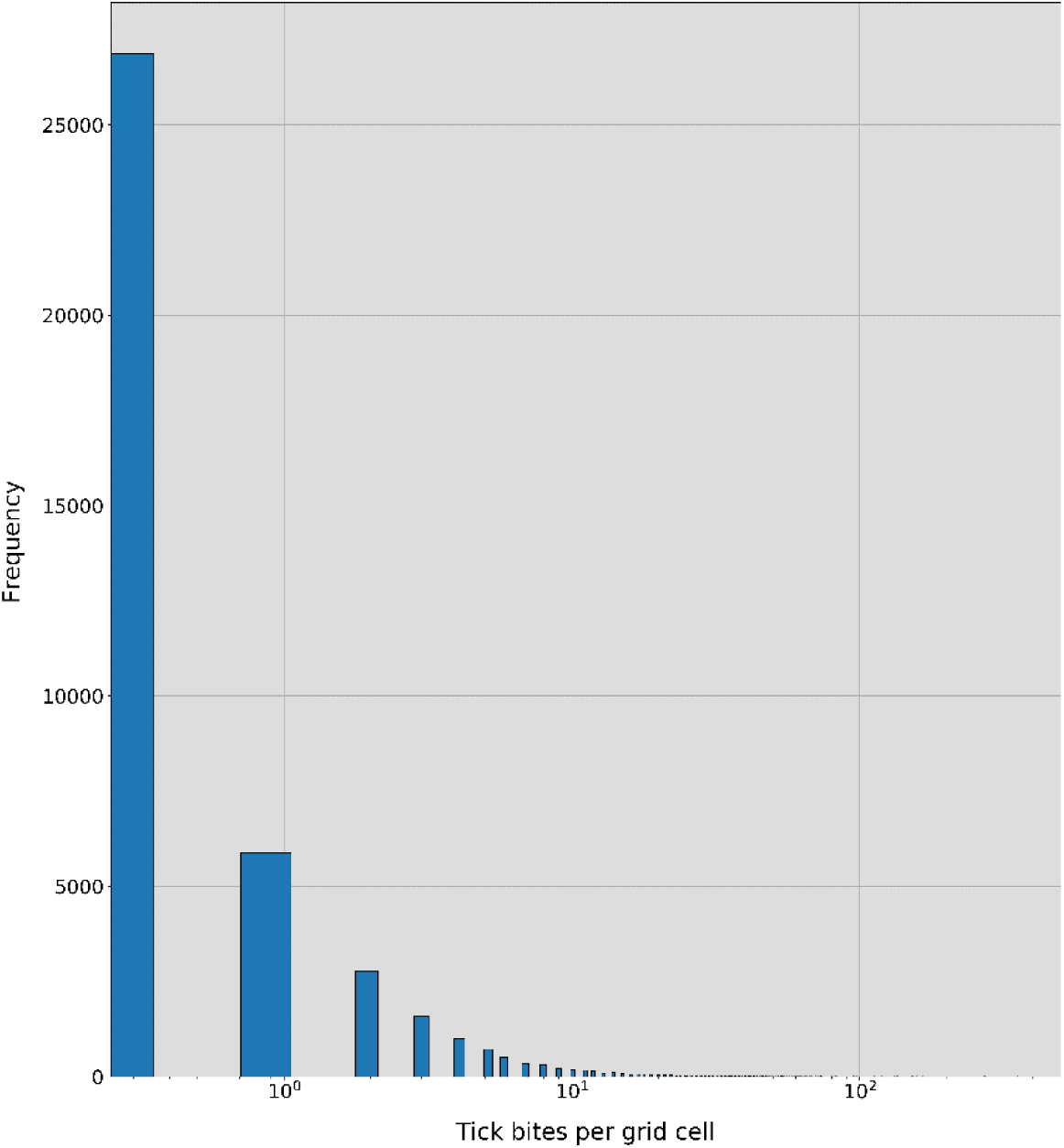
Histogram of the tick bites per grid cell. As seen, after the process of spatial aggregation described in Section 3.1, a skewed distribution with zero-inflation is created. Note that the X-axis is represented in log-scale to ease the visualization of the distribution. The number of grid cells containing more than 30 – 40 tick bite reports is almost negligible, but the distribution spans until a maximum of 353 tick bites per cell.

### 3.2 Ensemble learning from skewed and zero-inflated datasets

Random Forest (RF) (Breiman, 2001) is an ensemble learning method that can be used both for classifications an regression problems. The ensemble is formed by decision trees, whose individual predictions typically have a high variance, but when combined, they produce a robust and highly stable estimator (Louppe, Wehenkel, Sutera, & Geurts, 2013). RF combines bagging (Breiman, 1996) and the random subspace method (Ho, 1998). Bagging allows RF to see multiple variations of the input data and the random subspace method introduces randomness in the features presented to each tree during the learning phase. These two mechanisms are responsible of the diversity of the ensemble. RF predicts unseen samples by averaging the predictions of the trees in the ensemble.

RF and other canonical machine learning algorithms work under the assumption of having a similar number of samples per class or range. If this is not the case, the application of a canonical RF tend to produce results biased to the majority class or most common values (Japkowicz & Stephen, 2002; Krawczyk, 2016). Learning from a imbalanced (classification) or skewed (regression) dataset, is a non-trivial problem that started to receive attention in the early 2000s (He & Garcia, 2009). According to (Krawczyk, 2016) there are three categories of methods to learn from imbalanced or skewed data: 1) data-level methods; 2) algorithm-level methods; 3) hybrid methods. Data-level methods aim at balancing the dependent variable by applying over/under sampling techniques. Algorithm-level methods require the modification of the method in use to (partially) remove the bias towards the majority class or most common range of values. The hybrid methods combine the balancing of the dependent variable and a modification of the method in use. We propose an algorithm-level method to mitigate the effects of the data imbalance over the predictive power of RF.

As explained in section 3.1, the aggregation of the tick bite reports to create a 1km raster layer of tick bite risk created right-skewed and zero-inflated dataset (Figure 2). The sum of tick bites reported in each grid cell can be viewed as a discrete random variable that only takes non-negative values. This means that these reports can be modelled using well-known discrete probabilistic models for count data (hereinafter: count data models), such as Poisson (POI) and negative binomial (NB). Because of the large proportion of zero tick bites per grid cell, we also tested the zero-inflated versions of these models: the zero-inflated Poisson (ZIP) (Lambert, 1992), and the zero-inflated negative binomial (ZINB) (Greene, 1994) models. The difference between the original and the zero-inflated models is that in the latter type of models data is assumed to be derived from a two-stage process: 1) a Bernoulli trial deciding whether the event occurs or not (with probability *p*, the zero-inflation factor; 2) in case the event occurs, the counts will happen according to some rate λ. Note that this second process can also generate zeros. This two-stage process is convenient for the problem we are modelling. First, we check the presence of ticks (and humans) and if present, we check the “rate of transmission”, conceptually composed of visiting rates and biting rates.

Zero-inflated models have been used to predict TBE in a set-up in which the majority of the available samples had a zero (Stefanoff et al., 2018). However, this approach is limiting because count data models do not generally work well in set ups where there are complex non-linear interactions between the predictors and the response variable. In our work, the use of RF allows the inclusion of a wide array of predictors and the identification of the main ones to segment the problem into more homogeneous cases, which can afterwards be modelled using count data models. We propose a modelling approach that combines weak (i.e. decision trees) and strong (i.e. models for count data) estimators to improve the canonical form of RF. Figure 4 sketches our modelling approach where ensemble trees are only grown until their terminal leaves hold a minimal number of relatively homogeneous samples. These samples are subsequently analyzed with the four selected count data models (i.e. POI, NB, ZIP, ZINB). During the testing phase, each of the test samples will be propagated down each tree in the ensemble and will yield four predictions (one per count data model). The final prediction of the ensemble will be calculated by averaging the predictions of each model type, just like a canonical RF operates.

**Figure 3.**
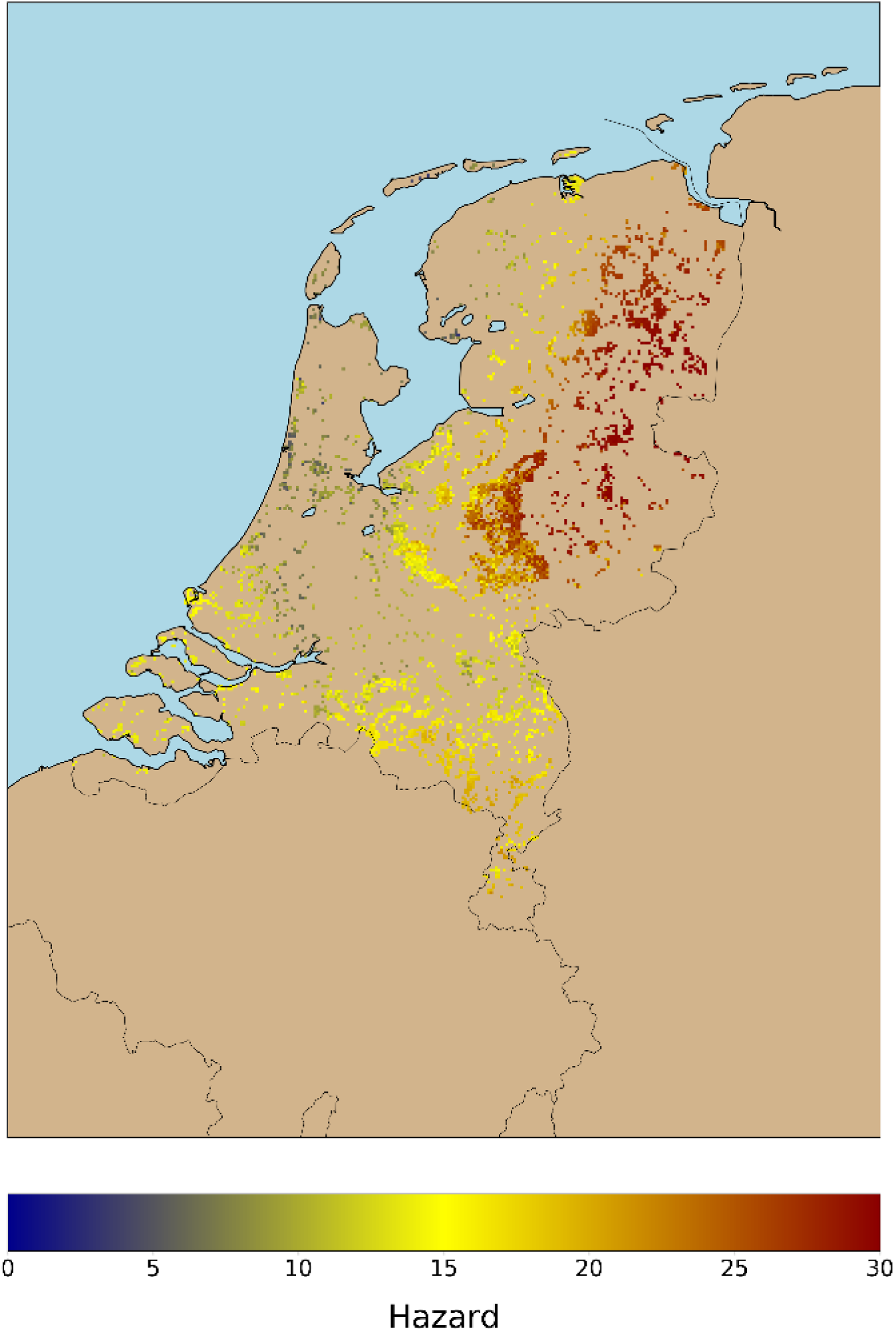
Tick activity per 1km grid cell. This tick activity estimation is provided by the data-driven model in (Garcia-Martí et al., 2017), which is capable of predicting daily tick activity. We run this model for each day during the period (2006-2014) and we calculated a robust long-term metric of hazard, showing the maximum mean tick activity for the entire period. As seen, hazard is minimum along the coastline and maximum in the northeast of the country.

**Figure 4.**
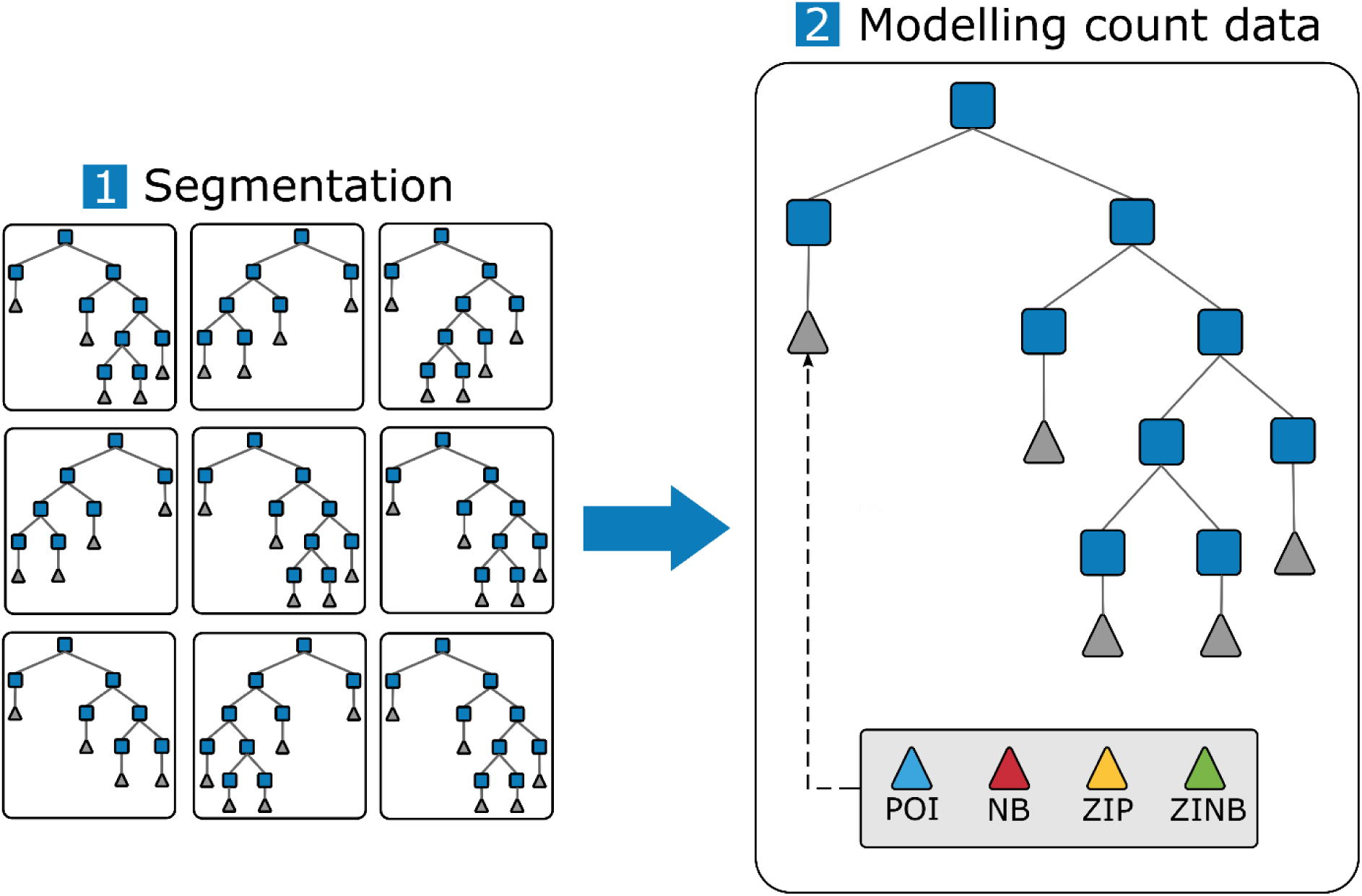
Proposed approach to couple RF and count data models (Poisson: POI, negative binomial: NB, zero-inflated Poisson: ZIP, and zero-inflated negative binomial: ZINB). First, the ensemble of decision trees is used to segment the samples into groups with similar characteristics. These trees are shallow trees, so that each of the leaf nodes contains a few hundred of samples. Second, we plug the selected count data models to each of the leaf nodes in the ensemble. The predictions yielded by the count data models are subsequently averaged to obtain the final prediction for each RF and count data model combination.

### 3.3 Modelling tick bite risk

The data and modelling approach described in the previous sections were used to model tick bite risk in the Netherlands. First, we explain the process of feature engineering applied to enrich each of the tick bite reports with hazard and exposure variables. Then, we describe our modelling experiments. Note that our work was developed using various Python libraries: numpy (Oliphant, 2006) to handle the multidimensional arrays, statsmodels (Seabold & Perktold, 2010) to fit the count data models, GDAL (GDAL Development Team, 2018) and Cartopy (Met Office UK, 2010) to process geospatial data and prepare the visualizations through map layers, matplotlib (Hunter, 2007) to prepare the rest of the figures, and SkillMetrics^1^ library and scipy (Oliphant, 2007) to obtain the statistical metrics used to evaluate the model.

#### 3.3.1 Feature engineering

In this study, we extend the ideas regarding human exposure described in (Zeimes et al., 2014). To do so, we use a substantial amount of official Dutch geospatial data, and of other user-contributed geo-sources, to derive accessibility and attractiveness metrics. Because of the aggregation of the tick bites to a uniform raster layer, the exposure metrics where calculated as the geographic distance between the center of each grid cell and a set of selected real-world features in which we expect the co-ocurrence of humans and ticks. As explained in Section 2.1, hazard metrics are extracted for forests and natural grasslands using the model developed by (Garcia-Martí et al., 2017). Table 1 presents the 19 exposure metrics and the 2 hazard metrics that were used to model tick bite risk in the Netherlands. The following paragraphs contain more details about each of the metrics.

**Table 1.**
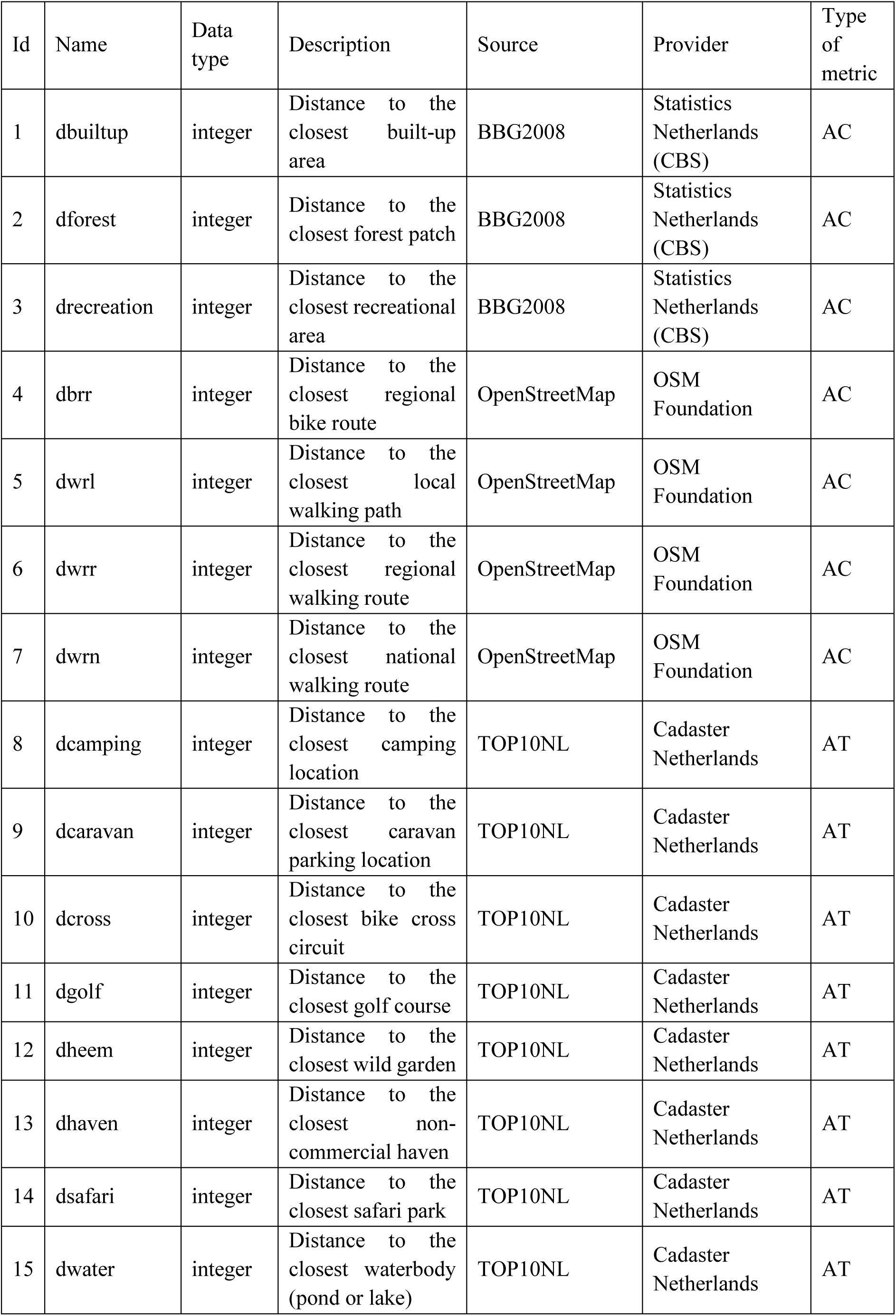
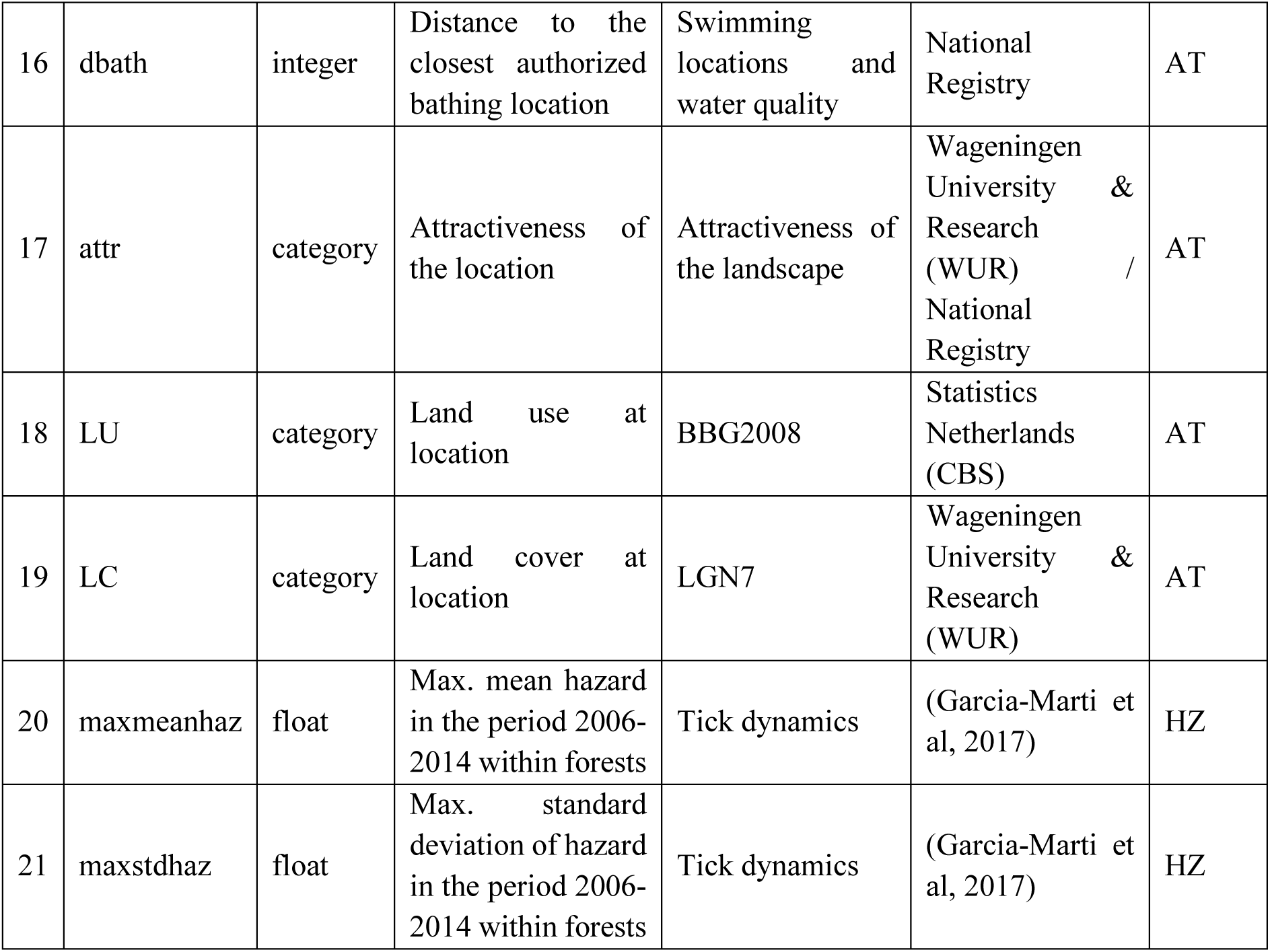
Features used in this work.

We derived accessibility metrics (Table 1, indices 1 – 7, Type AC) from the land-use map (BBG 2008) provided by Statistics Netherlands (CBS^2^), and from the transportation networks contributed by volunteers in OpenStreetMaps (OSM^3^). Using these data sources, we calculate the distance from each grid cell to a series of selected land uses and transportation networks. We compute the geographic distance (in meters) of each grid cell to the closest of selected BBG 2008 land use types, namely, forests, recreational areas and urban areas. We downloaded the latest snapshot of OSM for the Netherlands (last access, July 2018) and extracted the user-contributed cycling and walking networks. The former is available at local, regional and national scales whereas the latter is only available at the national scale. Note that the bike networks do not overlap so, for instance, the national routes do not include routes between small cities or forest patches, but longer routes connecting the edges of the country. We compute the geographic distance (in meters) between each grid cell and each of the selected cycling and walking networks.

We obtained attractiveness metrics (Table 1, indices 8 – 19, Type AT) by using data from the Dutch Cadaster, the Dutch National Registry, and WUR. From the Dutch Cadaster, we use the so-called functional polygons of their TOP10NL^4^ product. These polygons demarcate areas with 296 types of functions (Full list available: http://geoplaza.vu.nl/data/dataset/top10nl, last accessed November 2^nd^, 2018). Here, we selected 8 functions related to outdoor activities where humans could meet ticks: campings, caravan parks, bike cross circuit, golf courses, wild gardens, non-commercial havens, and safari parks. We also extract from the TOP10NL the location of all lakes and ponds in the country, since they can serve as attractors of visitors to nature due to its scenic beauty or recreational use. We include a publicly available map (Dutch National Registry) categorizing the attractiveness of the Dutch landscape (i.e. *Belevingswaarde van het landschap*^5^, last accessed July 5th, 2018) (Crommentuijn, Farjon, den Dekker, & van der Wulp, 2007; Roos-Klein Lankhorst, de Vries, Buijs, Bloemmen, & Schuiling, 2005). Finally, for each location, we extracted the land use and land cover categories from BBG 2008 and LGN7 database (produced by WUR), respectively.

The hazard metrics (Table 1, indices 20 – 21, Type HZ) are extracted from the model outlined in Section 2.1. This model, described in more detail in (Garcia-Martí et al., 2017), is based on nine years (2006-2014) of data collected by volunteers. These volunteers carried out a monthly sampling by cloth dragging of 15 vegetated locations in the Netherlands, counting the number of ticks per life stage (i.e. larvae, nymph, and adult). Our model was calibrated for the nymph life stage only, since nymphs pose the highest hazard to humans. The model also includes 101 biotic and abiotic environmental predictors. These predictors describe the tick habitat conditions (e.g. litter, moss), the occurrence of mast years for three tree species, weather conditions (e.g. temperature, evapotranspiration, relative humidity), satellite-derived vegetation indices (e.g. NDVI), and land cover. To account for the effect that short- and long-term weather conditions have on tick activity, we aggregated the weather data to 11 temporal resolutions (i.e. 1–7, 14, 30, 90, 365 days). We run our data-driven model for each day in the period 2006-2014. Then we computed the average of the maximum tick activity of each year and its standard deviation to obtain robust and long-term proxies for tick hazard in forests and natural grasslands locations. This means that outside of these locations, the hazard is unknown. In this case, we are unable to use value imputation, since this would require imputing values for most of the country. Instead, outside of these locations, we used a symbolic value of minus one. This value is meant to separate locations for which we have and do not have hazard data. The selection of this value is backed up by recent research (Heylen et al., 2019), which shows that tick densities decrease along the forest-urban land use transition. Thus, the symbolic value of minus one helps at grouping together samples without hazard and samples with low hazard, which tend to occur outside forests.

#### 3.3.2 Experiments

The spatial aggregation and feature engineering described in sections 3.1 and 3.3.1, resulted in a matrix with 36,866 rows and 21 columns. Each row represents a grid cell and each column the E or H features selected for this work. A series of experiments were designed to identify a tick bite risk model that can handle the skewness and zero-inflation present in this matrix. First, we randomly selected 60% of the data for training all the models and reserved the remaining 40% for testing them. Then, we defined a range of values for the two main RF parameters of our ensemble: 1) the number of tree estimators; and 2) the number of samples per terminal leaf node. We trained ensembles using 10, 20, and 50 trees and where each tree had at least 100, 200, 400, 600 and 800 samples per terminal leaf. The number of samples per leaf node determines the “level of development” of the trees in the ensemble. Thus, experiments with few samples per leaf node (e.g. 100 samples) create deep trees close to full development, whereas shallow trees are created when there are many samples per leaf node (e.g. 800 samples). In total, 15 RF ensembles were trained using the same split of training and test samples. Subsequently, these RF ensembles were crossed with the four discrete probability models for count data (i.e. POI, NB, ZIP, ZINB), which were fitted using a non-parametric approach (i.e. without having to specify any hyper parameter), using a Nelder-Meade optimization routine to obtain the maximum likelihood estimates of the parameters of the distributions.

Two issues could hamper the fitting of the count data models: excessive skewness or excessive zero-inflation. The selected count data models can deal with skewed distributions, but the segmentation carried out by RF might leave the leave nodes with a subset of samples highly skewed towards zero (i.e. 85% - 100% of zeros). We explored how often these circumstances occur for each tree in the ensemble and we found out that in average, the fitting does not converge in 5% - 9% of the leaf nodes in the ensemble. In those cases, we keep the default behavior of a canonical RF, which is returning the mean of the samples falling in that node. Finally, model performance is checked with the test dataset. For this, we track the itinerary of each of the test samples down the tree, and identify the leaf node in which it ends up. Then, we pass this sample to each of count data models fitted with data from that node to get the prediction of that tree. We do this for each tree in the ensemble, and we average these tree-based predictions to get the final (ensemble) prediction, following the default behavior of canonical RF models.

A modified Taylor diagram is used to graphically summarize the test results. This diagram shows three statistics in a single plot: the standard deviation, the root-mean-squared deviation (RMSD), and Pearson’s correlation coefficient. Taylor diagrams represent the relationship between the three variables, which essentially lie on a 2D manifold and are projected onto a 2D flat geometry without loss of information. We use this diagram because an accuracy metric like RMSD alone is not informative in the case of heavily skewed data, where also measures of dispersion play an important role. Since our distribution is skewed and zero-inflated, we substituted Pearson’s coefficient by Spearman and Kendall Tau ranked correlation coefficients. The ensemble and count data models combination that yield reasonable trade-off between RMSD, standard deviation and correlation coefficient are then used to create tick bite risk maps for the Netherlands. The map created when using a canonical RF is added to the list of best models to be able to evaluate the advantages of our approach. Finally, we intersect these maps with the human exposure layers showed in Figure 5.

**Figure 5.**
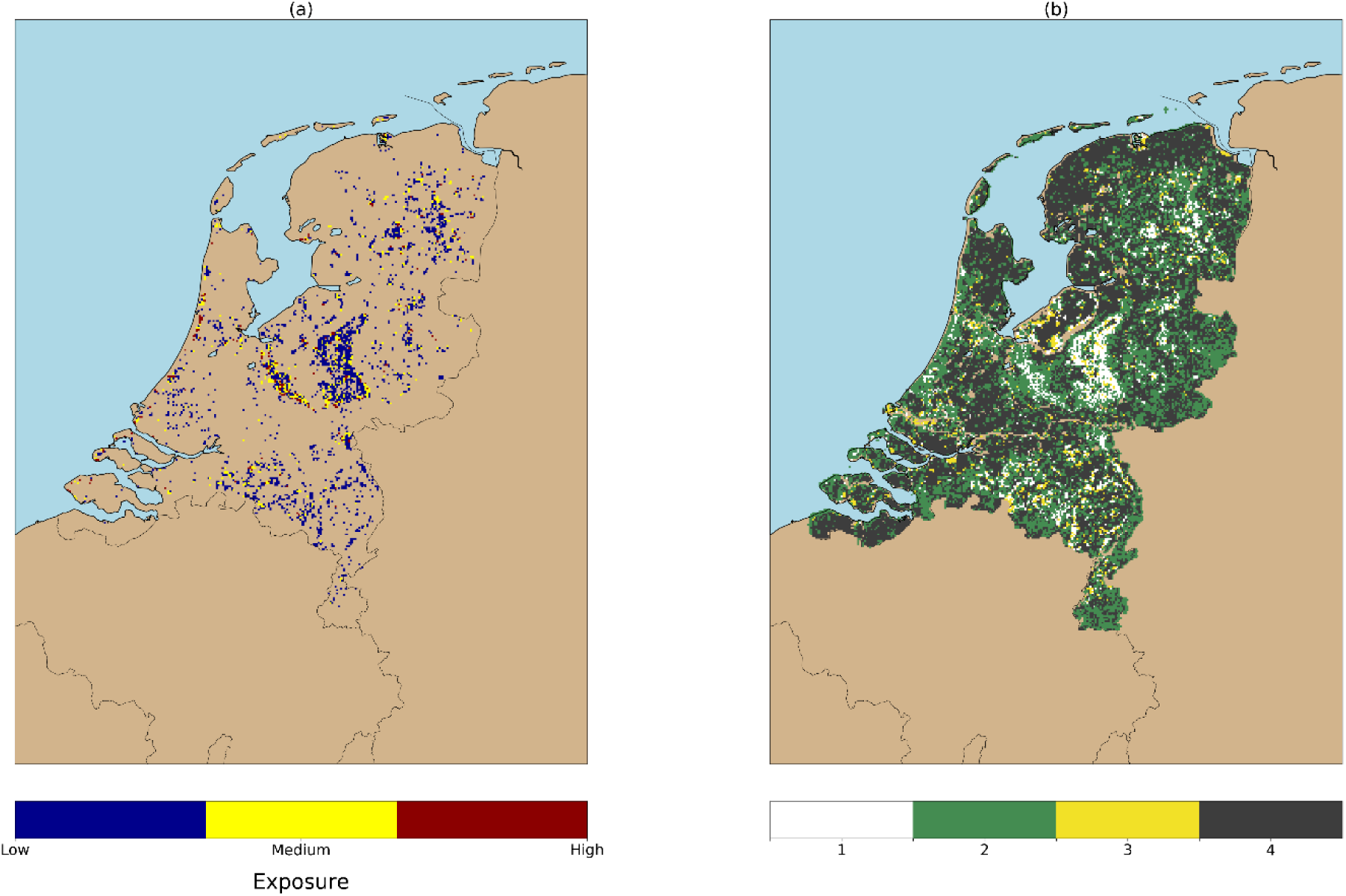
(a) Human exposure to tick bites classified in three categories. (b) Class 1 in this map correspond to the classes from (a), class 2 represents tick bites reported outside forests, class 3 represents forests with no tick bites recorded, and class 4 shows locations where no tick bites were reported during the study period. These results can be found in (Garcia-Marti et al., 2018), and we cross them with the tick bite risk maps obtained in this work to explore the risk per human exposure category (Figure 9).

## 4 Results

Figure 6 shows the ability of the four count data models and the canonical RF to fit the skewed distribution of tick bites in the different set-ups. Overdispersion is better fitted by POI and ZIP models compared to NB and ZINB, since the former models yield values between 0 and 30 tick bites per grid cell, whereas NB and ZINB are barely able to predict beyond 10 tick bites per cell. Interestingly, the zero-inflation seems to be better captured by NB and ZINB than POI and ZIP, as seen by the frequency of predicted zeros of these models is similar to the original distribution. RF performs similarly in all the prepared set-ups, and seems unable to predict over a wide range of values, most values typically being constrained to below 5 tick bites. In general terms, the NB, ZIP, and ZINB models seem to capture reasonably well the original distribution, but the POI model and the canonical RF do not perform well: the POI model yields predictions with a frequency considerably higher than the original values, whereas RF is unable to predict beyond few tick bites per grid cell. In addition, POI and RF are not able to capture the zero-inflation. As seen, the predicted distributions do not seem to considerably improve or deteriorate based on the increasing number of samples per leaf node (i.e. 100-600 samples), but the experiments with shallow trees (i.e. 800 samples) seem to have a negative impact in the ability of the models to predict zeros.

**Figure 6.**
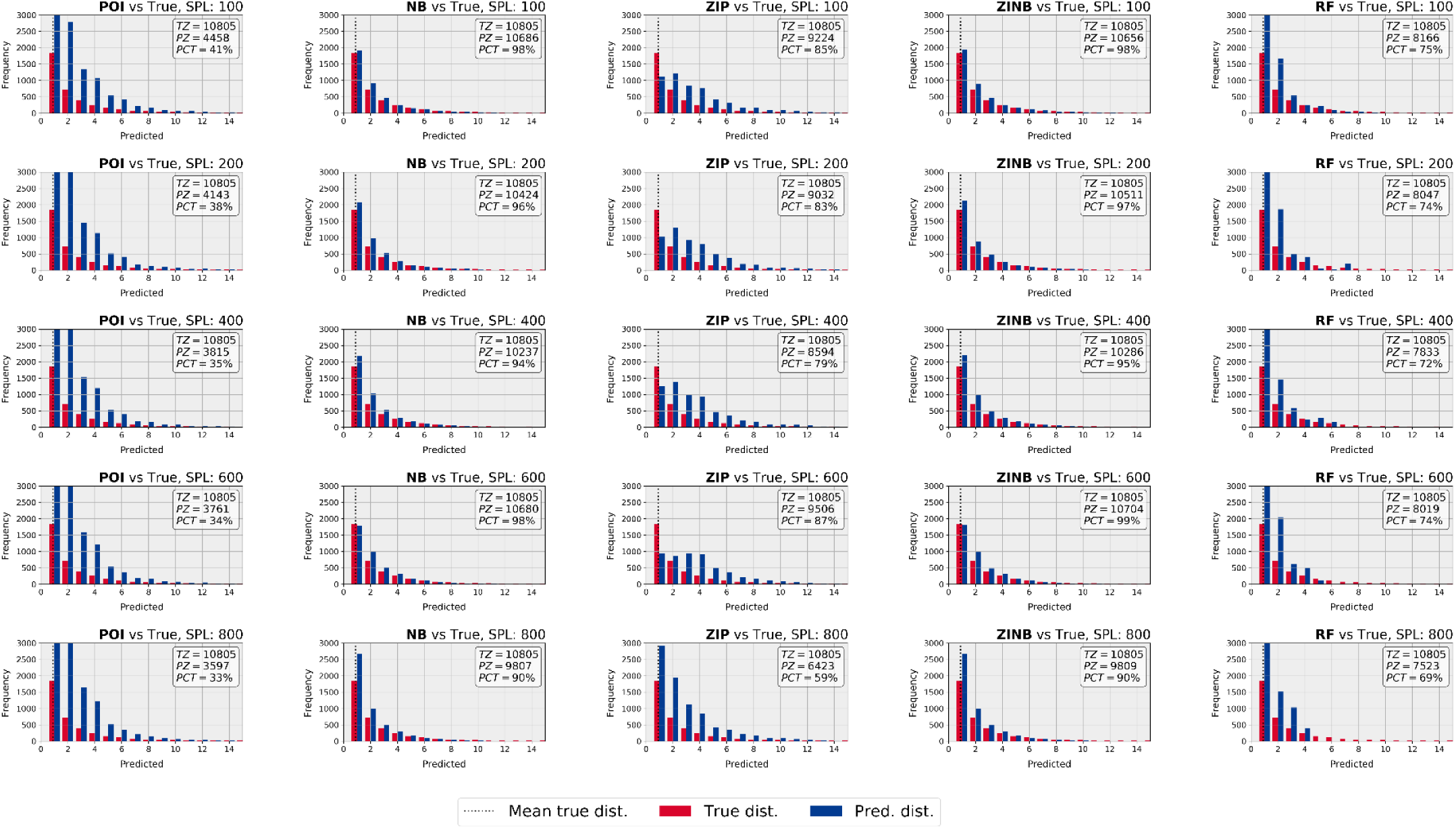
Histograms of original (red) and predicted (blue) distributions for an ensemble with 20 trees and an increasing number of samples per leaf node. Note that for visualization purposes, the axes have been limited and the zeros are summarized in the text box within each subplot, thus showing the number of true zeros, predicted zeros and the associated percentage. The first four columns correspond to the count data models, whereas the last column shows the performance of the canonical RF. As seen, POI and ZIP can predict for a wider range of values, whereas NB and ZINB are good at predicting zeros and the low part of the distribution.

Figure 7 shows the performance of the ensemble in two modified Taylor diagrams. Each of the colored symbols represents an ensemble with a concrete number of tree estimators (T) and samples per leaf node (S). A visual inspection of the diagrams reveals that all ensembles yield predictions that are strongly correlated (i.e. correlation > 0.8 for Spearman’s and > 0.7 for Kendall’s Tau) with the tick bite data. The Taylor diagrams also show that the RMSD of these models is within a reasonable and stable range (i.e. 1 – 6) for all the experiments. However, in this work we are not only interested in models with a high correlation and low error, but also in those providing a realistic range of predictions, which is given by the standard deviation (stdev) represented by the dotted radial axes. The models present a variable skill to account for overdispersion (i.e. stdev 1 – 8). Considering these three statistical metrics, we think that the models better performing are located under the arc created by RMSD=2. Using the pink hexagon as a reference point, we can see that there are NB, ZIP, and ZINB models below this arc present a high Pearson/Kendall correlation (i.e. >0.9), a low RMSD (i.e. <2) and a fair range of stdev (i.e. 2 – 5). Out of these selected models, we can see that 2 ZIP and 1 ZINB models present a higher skill to model overdispersion (stdev > 4), whereas the small cluster of NB and ZINB models under the arc are better suited to predict zero-inflation. These diagrams also show that the optimal experiments correspond to 200-600 S and 20-50 T. To create the tick bite risk maps, we select the experiments with 200 samples per leaf node and 20 tree estimators since we believe they provide the best results. Figure 8 shows the tick bite risk maps produced by the four count data models (a-d), by the canonical RF (e), and a zoom-up of the maps obtained with the ZIP and ZINB models (f-g). The application of POI and ZIP models at the country level create maps that present a wide range of predicted tick bites. The NB and ZINB models yield maps that are visually less prominent, and the predictions of RF are mostly uniform throughout the territory and do not show any remarkable pattern. In Figure 6 and 7 we see that NB and ZINB present a higher skill to model zero-inflation, which means that they perform better than POI or ZIP at delineating regions with a low tick bite risk. In this sense, NB and ZIP mark large areas in the northwest (e.g. province of Friesland) and in the center west (e.g. region of the Groene Hart) of the country, either as very low or inexistent risk of tick bites. Figure 6 and 7 also show that ZIP has a better ability to predict over dispersed data, which is particularly suitable to identify less prominent risky locations, such as the patchy forest structures of the northeast of the country (e.g. provinces of Groningen and Drenthe) and the forests in the south (e.g. province of Noord-Brabant). The inspection of the zoomed ZIP and ZINB models (Figure 8; f-g) shows that the risk is maximum in popular recreational locations. The Veluwe national park, the Utrechtse Heuvelrug forests, and the recreational areas along the coast present the highest tick bite risk of the country.

**Figure 7.**
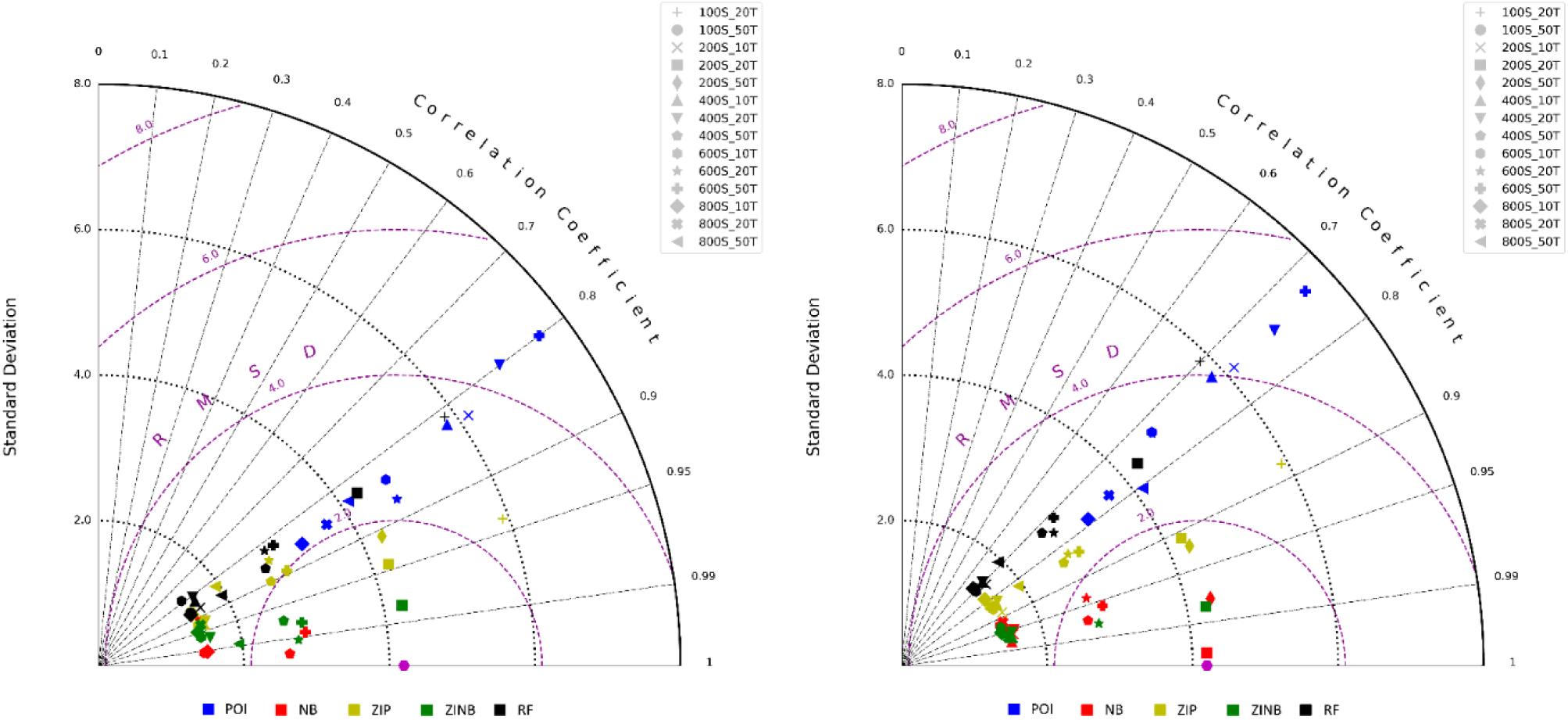
Modified Taylor diagrams showing the performance of the models based in three metrics: standard deviation, RMSD, and a correlation coefficient (left: Spearman’s, right: Kendall Tau), which are represented by the Y, circular, and radial axes, respectively. Each of the colored symbols represents an ensemble with a concrete number of tree estimators (T) and samples per leaf node (S). The models better performing are located under the arc created by RMSD=2, since they present a high Pearson/Kendall coefficient, low RMSD, and a fair standard deviation. Out of these selected models, we can see that 2 ZIP and 1 ZINB models present a higher skill to model overdispersion (i.e. std. dev.>4), whereas the small cluster of NB and ZINB models under the arc are better suited to predict zero-inflation. As seen, experiments with in the range of 200-600 samples per leaf node seem to perform optimally in both diagrams.

**Figure 8.**
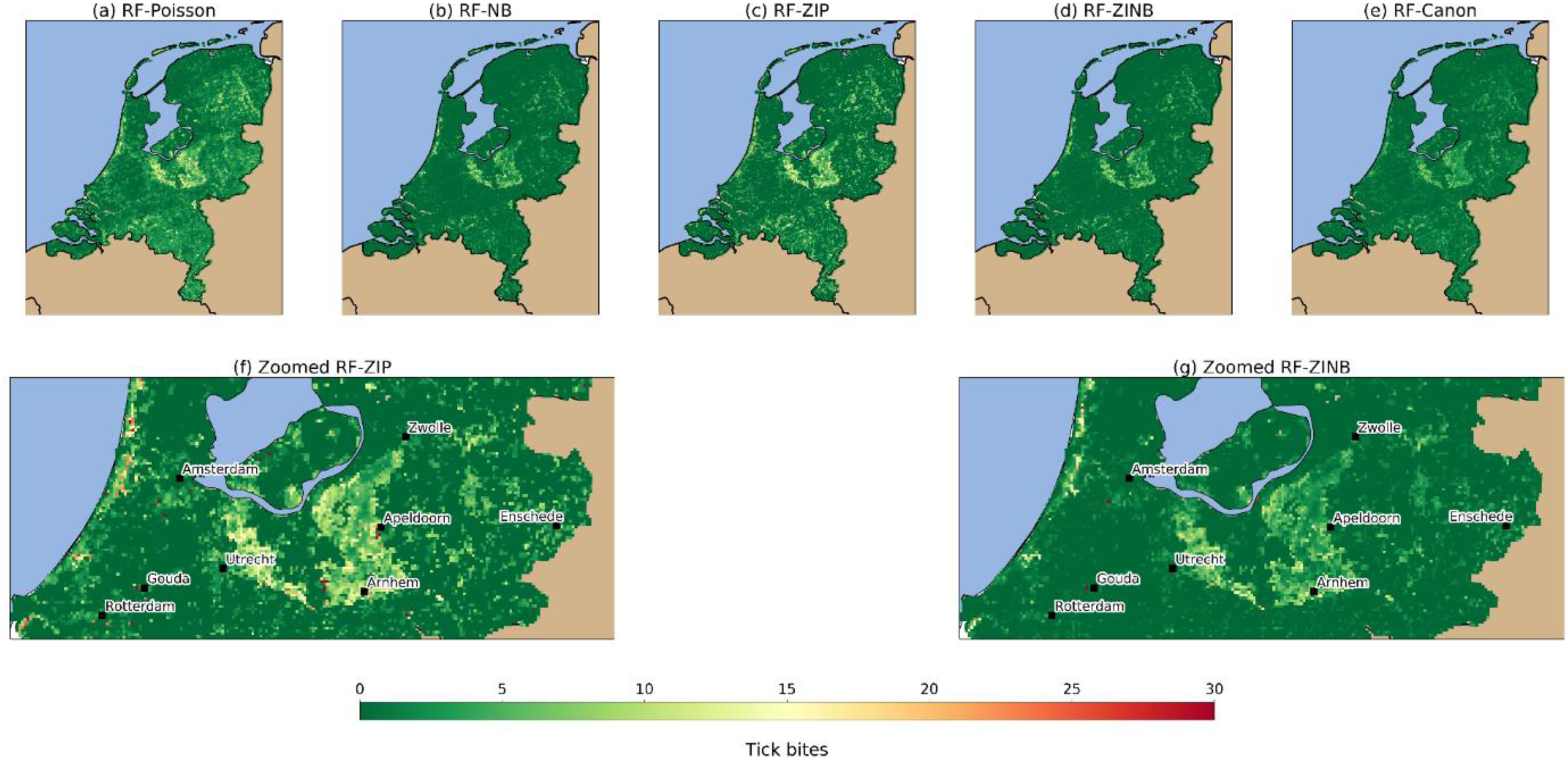
Tick bite risk maps produced by combining RF with each of the count data models. The upper row (a-e) shows the general overview of the models, whereas the bottom row shows a close-up for the ZIP (f) and ZINB (g) models. The NB (b) and ZINB (d) models are better suited to delineate regions with low or inexistent tick bite risk, thus they present sharper declines between different land uses. The ZIP (c) model is capable of predicting the risk of tick bite over a range of values, this is why its associated map presents smooth and gradual changes across the study area. POI (a) and RF (e) are over/under predicting, respectively, since the former finds risk in most locations of the country, whereas the latter yields and almost-homogeneous prediction. The visual inspection of the zoomed models (f, g) identify popular places for recreation intensely visited by citizens. The forested areas between Utrecht, Apeldoorn, and Arnhem, together with the recreational areas along the coast are regions where tick bite risk is particularly high.

Figure 9 depicts the risk of tick bite classified by the exposure levels found in Figure 5. We show a plot for each count model (a-d), for the canonical RF (e), and the original volunteered reports from NK and TR (f) classified by type of human exposure as well. The models have different skills to predict risk for each of the exposure classes in Figure 5. Considering the low, medium, and high exposure classes in Figure 5a, we see that ZIP better captures the overdispersion of data, since its interquartile range across classes (i.e. 0 – 10 TB/cell) is longer than NB and ZINB models (i.e. 0 – 6 TB/cell). Also, NB and ZINB present a higher skill at modelling the low exposure class, since it coincides with the original tick bite distribution (i.e. 0 – 4 TB/cell). ZIP provides more flexibility at predicting for the medium (i.e. 3 – 9 TB/cell) and high (i.e. 3 – 10 TB/cell) exposure classes, since these they span a range resembling the original distribution (i.e. 0 – 6 TB/cell and 0 – 14 TB/cell, respectively). Regarding the exposure classes in Figure 5b, we can see that ZIP seems to capture well the category of tick bite risk in non-intensively visited forests. The NB, ZIP, and ZINB models are not able to predict a range for the category of tick bites outside forests. The canonical RF shows a poor performance across the exposure classes, since it is only able to predict for a narrow margin of the original distribution. Based in the results provided in this section, we believe that, overall, the ZIP and ZINB models present stable predictions and the ability of modelling overdispersion and zero-inflation, respectively.

**Figure 9.**
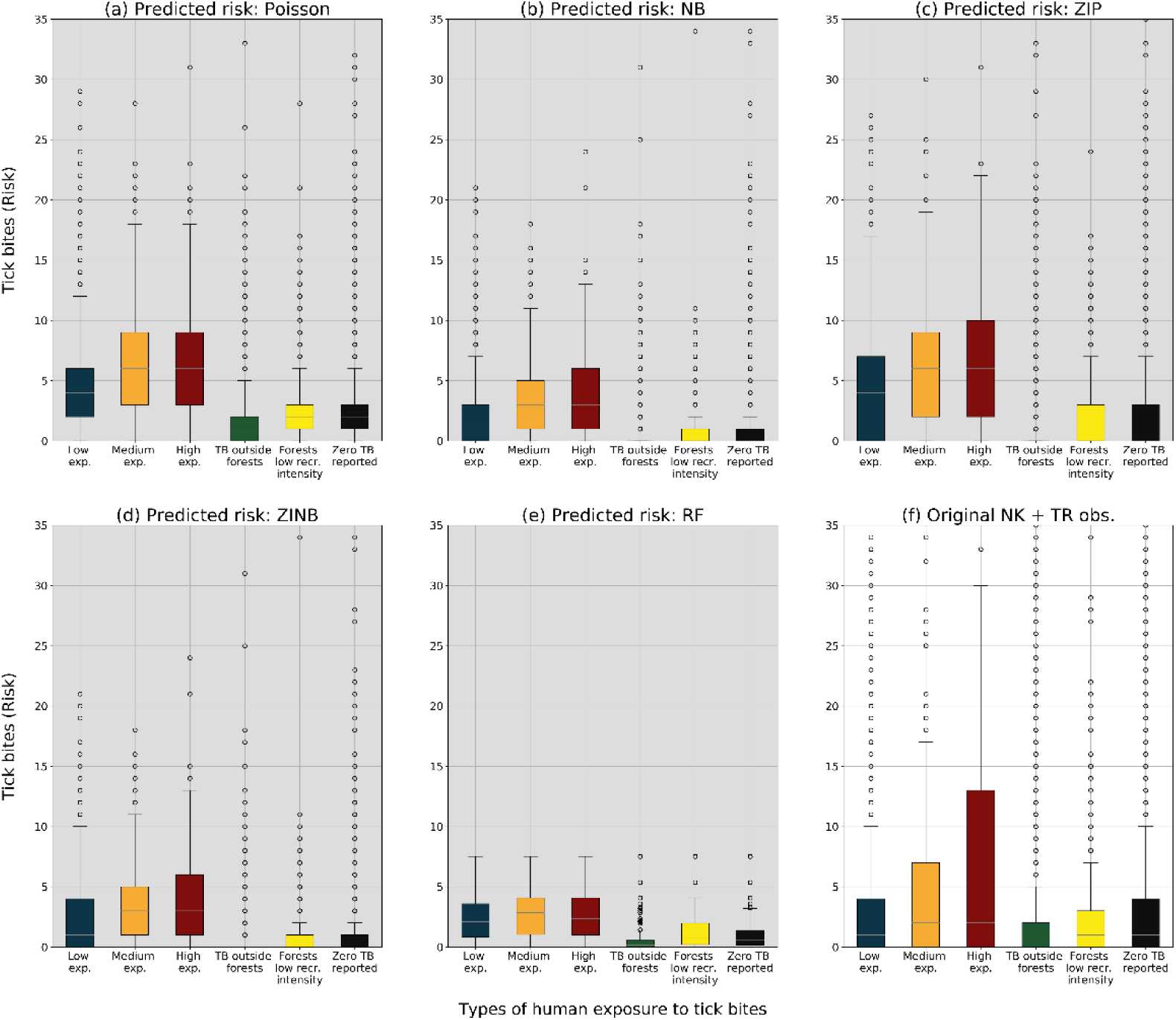
Tick bite risk classified based on the human exposure classes from Figure 5. The subplots show the predicted distributions per human exposure class for each of the count data models (a-d) and RF (e), and the type of human exposure using the original volunteered observations from NK and TR (f). The models present a different skill at predicting for each of the exposure classes. For example, ZIP and ZINB yield predictions for the low exposure class very similar to the original ones, whereas ZIP has analogous predictive capabilities for the forests with a low recreational intensity. Note, however, that although ZIP and ZINB can model the medium exposure class reasonably, none of the used models are able to capture the high skewness present in the high exposure class. The canonical RF is not able to predict the over dispersion of the original dataset.

## 5 Discussion

In this work we illustrate that canonical RF models have difficulties capturing skewed distributions and we present our approach conceived to mitigate the effects of biasing the model towards the mean. To do so, we apply an algorithm-level modification of RF (Krawczyk, 2016), by combining weak estimators (i.e. decision trees) with robust estimators (i.e. count data models). By doing this, we keep two important characteristics of both types of estimators: a fast segmentation of the samples, and a realistic prediction of the tick bite risk. Thus, the integration of the segmentation capabilities of RF and the count data models creates a robust combined estimator.

Due to the skewed and zero-inflated nature of the tick bites per grid cell, our work does not aim at creating a model with the lowest performance metrics (i.e. low RMSD and standard deviation), but a model that finds the trade-off between the error and the capability of predicting tick bites over the reaslistic range of data values. For this, we tested various model configurations. The metrics represented in Figure 7 show the performance of the models based in three metrics: the standard deviation, the RMSD, and a correlation coefficient. Based on these three metrics, we think the that the ZIP and ZINB models are the ones performing better, since they present good correlation coefficients, reduced RMSD and are able to capture overdispersion or zero-inflation, respectively, which can open the door to multiple applications in ecological modelling and public health.

The presented maps illustrate that the proposed approach can be used to estimate the tick bite risk in a location. The NB and ZINB models seem adequate for low-risk detection, since they perform better with zero-inflation in data, which subsequently enables the identification of low-risk regions. Then, the ZIP models are more suitable to fit over dispersed data, which enables the quantification of the risk within a wide range of values. Visually, this means that NB and ZINB maps identifies large regions with low-risk with sharp declines between different land uses, whereas the predictions yielded by ZIP show a richer range of predictions that can help at location risky locations in the country. The selected ZIP and ZINB models are able to identify locations of high risk in popular recreational places (e.g. Veluwe national park, coastal recreational areas), but they also have proved useful at detecting risky locations which are less intensively visited by citizens (e.g. patchy forests in the provinces of Noord-Braband, Drenthe, and Groningen). We believe that these maps can support several public health interventions intended to decrease the number of tick bites.

Using the categories from Figure 5 and the map layers in Figure 8, we inspected the risk of tick bites in function of human exposure inside and outside forests. In Figure 9 we see that some of the models are able to predict reasonably well for certain human exposure categories. For example, ZIP and ZINB yield predictions for the low exposure class very similar to the original ones, whereas ZIP has analogous predictive capabilities for the forests with a low recreational intensity. Note, however, that although ZIP and ZINB can model the medium exposure class reasonably, none of the used models are able to capture the high skewness present in the high exposure class. This limitation suggests that human exposure in highly visited locations might need additional features to better characterize the human activities outdoors. Considering all insights together, we think that these results suggest that a combination of RF and ZIP would be the most suitable one to estimate the tick bite risk in a location, whereas the combination of RF and ZINB would be adequate to detect locations with zero or very low risk.

In this work we encountered four hurdles. First, finding a proper validation metric for skewed distributions was challenging, because the most commonly used measures of model performance use statistical measures of location, not of dispersion, whereas in this case we are equally interested in predicting the dispersion. The (modified) Taylor diagram can help at evaluating the models because it can represent three statistical metrics in a single chart. Second, the TB collection is self-reported by each user of NK and TR. This means that this is a source of spatial inaccuracy based on the level of map literacy and spatial awareness of each user. With the current data collection, we are not able to quantify, nor correct, for this spatial inaccuracy. This means that at the time of the feature engineering we might be characterizing an observation which is incorrectly placed. We acknowledge the importance of citizen science campaigns, but we recommend that further data collection campaigns dedicate effort to find the positional accuracy of each observation. Third, there is a small fraction of the non-parametric count data model fittings that fail to converge due to excessive data imbalance for the optimization routines. Further work should aim at incorporating statistical knowledge to improve the fitting procedure, so that all models converge and contribute to the joint prediction of the ensemble. Fourth, the hazard model used in this work can produce a robust estimation of tick activity within forests, but not on other land uses. Thus, in this work the contribution of E and H could only be estimated in forested areas, whereas in the remaining land uses the model is entirely driven by E features. Further work should aim at combining different hazard metrics (e.g. tick suitability) to obtain a continuous picture of tick hazard throughout the country. This improved hazard metrics could help at disentangling which of the two factors (i.e. E or H) is dominant for each location, and thus would allow a deeper understanding of the factors of tick bite risk.

## 6 Conclusion

In this work we illustrate how canonical machine learning algorithms like RF may not perform well at modelling a skewed and zero-inflated distribution, and we present our algorithm-level solution to mitigate the bias towards the mean. Our approach consists in modifying the default behavior of RF by combining weak estimators (i.e. decision trees) with robust estimators (i.e. count data models). Thus, we connect four discrete probability models for count data (i.e. Poisson, negative binomial, zero-inflated Poisson, and zero-inflated negative-binomial) to each of the leaf nodes of RF. Subsequently, we enable RF to predict for skewed and zero-inflated distributions, which constitutes a methodological step forward in the machine learning field. We used this modified RF to model tick bite risk using volunteered reports collected by two Dutch citizen science projects. We extend the current state of the art on tick bite risk modelling by devising and integrating a wide array of hazard and exposure metrics. By doing this, we are able to create tick bite risk maps for the Netherlands, and to explore the risk based on human exposure. We hope that this double contribution can help other researchers across multiple fields at modelling skewed and zero-inflated distributions using machine learning methods. Finally, we believe that this work also demonstrates that the incorporation of volunteered data to a scientific workflow is not only possible, but recommended to model fine-grained phenomena that escape classic monitoring networks.

## Acknowledgements

The authors would like to thank Dr. A.J.H. van Vliet (Wageningen University) for providing access to data on tick bites from Natuurkalender and Tekenradar. The authors also would like to thank Dr. C.C. van den Wijngaard (RIVM), and M. Harms (RIVM) for co-providing the Tekenradar dataset. Last but not least, we would like to thank the anonymous citizens that contribute to the Natuurkalender and Tekenradar platforms.

## Footnotes

1 https://github.com/PeterRochford/SkillMetrics

2 https://www.cbs.nl/en-gb

3 https://www.openstreetmap.org

4 https://zakelijk.kadaster.nl/-/top10nl

5 https://data.overheid.nl/data/dataset/49505-belevingswaarde-van-het-landschap

